# Matching Variants for functional characterization of genetic variants

**DOI:** 10.1101/2023.02.22.529565

**Authors:** Sebiha Cevik, Pei Zhao, Atiyye Zorluer, Wenyin Bian, Oktay I. Kaplan

**Affiliations:** Rare Disease Laboratory, School of Life and Natural Sciences, Abdullah Gul University, Kayseri, Turkey; School of Applied Science and Engineering, Fuzhou Institute of Technology, Fuzhou 350014, China; SunyBiotech Co., Ltd, Fuzhou 35000, China

## Abstract

Rapid and low-cost sequencing, as well as computer analysis, have facilitated the diagnosis of many genetic diseases, resulting in a substantial rise in the number of disease-associated genes. However, genetic diagnosis of many disorders remains problematic due to the lack of interpretation for many genetic variants, especially missenses, the infeasibility of high-throughput experiments on mammals, and the shortcomings of computational prediction technologies. Additionally, the available mutant databases are not well-utilized. Toward this end, we used *Caenorhabditis elegans* mutant resources to delineate the functions of eight missense variants (V444I, V517D, E610K, L732F, E817K, H873P, R1105K, and G1205E) and two stop codons (W937stop and Q1434stop), including several matching variants (MatchVar) with human in ciliopathy associated IFT-140 (also called CHE-11)//IFT140 (intraflagellar transport protein 140). Moreover, MatchVars carrying *C. elegans* mutants, including IFT-140(G680S) and IFT-140(P702A) for the human (G704S) (dbSNP: rs150745099) and P726A (dbSNP: rs1057518064 and a conflicting variation) were created using CRISPR/Cas9. IFT140 is a key component of IFT complex A (IFT-A), which is involved in the retrograde transport of IFT along cilia and the entrance of G protein-coupled receptors (GPCRs) into cilia. Functional analysis of all ten variants revealed that P702A and W937stop, but not others phenocopied the ciliary phenotypes (short cilia, IFT accumulations, mislocalization of membrane proteins, and cilia entry of non-ciliary proteins) of the IFT-140 null mutant, indicating that both P702A and W937stop are phenotypic in *C. elegans*. Our functional data offered experimental support for interpreting human variants, by using ready-to-use mutants carrying MatchVars and generating MatchVars with CRISPR/Cas9.

## Introduction

Rare diseases are classified as disorders that affect a small number of people. A recent estimation revealed that 3.5–5.9% of the world’s population will be affected by nearly 6200 different rare diseases, 72% of which have genetic foundations (Nguengang Wakap et al., 2020). The genetic diagnosis of many rare diseases was hindered due to a lack of extensive human sequencing data that would have provided allele frequency estimates for each rare human variant. Because it is thought that rare genetic diseases are more likely to be caused by genetic variants with rare representation in human populations, researchers and clinical scientists sequenced a small sample of individuals with and without diseases to estimate the prevalence of a disease-causing variant. To help diagnosis of genetic diseases, the large genome consortiums, including the 1000 Genomes Project, Exome Sequencing Project (ESP), the Exome Aggregation Consortium (ExAC), the genome aggregation database (gnomAD), and Trans-Omics for Precision Medicine (TOPMed) Program undertook the steps necessary to create the human variation databases that could provide the reference allele frequency (Exome Aggregation Consortium et al., 2016; Fu et al., 2013; Karczewski et al., 2020; Taliun et al., 2021; The 1000 Genomes Project Consortium, 2010). Despite substantial technological advancements in genomic sequencing and the availability of great databases for the reference allele frequency, the analysis of genomics data, the interpretation of each variant for that particular type of disease still presents a major challenge for patients and their families in the diagnosis of rare diseases. Because large-scale human genome sequencing has revealed that there are a significant number of genetic variations (705,486,649 variants) in human populations. Furthermore, the underrepresentation of different ethnicities and populations in gnomAD and TOPMed makes it difficult to classify missing variants (Karczewski et al., 2020; Taliun et al., 2021). For this reason, a large percentage of human variants are classified as “variant of uncertain significance” (VUS) (Pir et al., 2022).

Many computational prediction tools are now available for predicting the functional impact of genetic variants, but the classification of missense variations does not always correspond with the development of the disease (Adzhubei et al., 2010; Chun and Fay, 2009; Davydov et al., 2010; Kircher et al., 2014; Ng, 2003; Reva et al., 2011; Schwarz et al., 2014; Shihab et al., 2013). For instance, employing four different pathogenic prediction technologies, including Align-GVGD, SIFT, MutationTaster2, and PolyPhen-2, to classify 670 VUS in *BRCA1* and *BRCA2* revealed the insufficiency of these tools for pathogenicity prediction (Ernst et al., 2018). Furthermore, high-throughput experiments have emerged as an alternate method of classifying variants, but scaling them to scan millions of variants is currently not feasible (Findlay et al., 2018; Giacomelli et al., 2018; Glazer et al., 2020).

Model organisms serve as a valuable testing system to discover the functional impacts of human genetic variants. Indeed, the ease of implementation of CRISPR/Cas9 genome editing has paved the way for such analysis, aiding to classify variants into functional categories, including phenotypic/pathogenic variants (McDiarmid et al., 2018; Wong et al., 2019). Furthermore, we recently unveiled a matching (equivalent) variants (MatchVars) search engine for *C. elegans*, mice, and human variants (Pir et al., 2022). Analysis of the Australian Phenome Bank (APB) and the Million Mutation Project databases revealed that they produced many mutants with missense variants, including MatchVars, whose potential had previously received less attention.

In the current study, we took advantage of available *C. elegans* mutants from Million Mutation Project database and concentrated on mutants bearing variants in IFT140 (intraflagellar transport 140). Intraflagellar transport (IFT) is a cilia-specific bi-directional transport activity that has been preserved throughout evolution and is essential for the formation of cilia (Blacque, 2008; Kozminski et al., 1993). IFT140 is required for the return of IFT complex from the ciliary tip to the ciliary base. Furthermore, IFT140 is found to regulate the ciliary entry of G protein-coupled receptors (GPCRs) (Absalon et al., 2008; Mukhopadhyay et al., 2010). *IFT140* was implicated in several ciliopathies, including Short-Rib Thoracic Dysplasia 9 with or Without Polydactyly (SRTD9: 266920), [also known as Mainzer-Saldino syndrome (MSS), conorenal syndrome and Jeune asphyxiating thoracic dystrophy (JATD)], nonsyndromic retinitis pigmentosa, syndromic congenital retinal dystrophy, and autosomal dominant polycystic kidney disease (ADPKD) (Ali et al., 2023; Bayat et al., 2017; Bifari et al., 2016; Helm et al., 2017; Schmidts et al., 2013; Xu et al., 2015). However, the majority of variants (80%) in human *IFT140* submitted to ClinVar are categorized as VUS (Landrum et al., 2018). We first identified MatchVars between *C. elegans* IFT-140 and human IFT140 using ConVarT, then we obtained 12 mutants harboring missense, including MatchVars and stop codons (V444I, V517D, E610K, L732F, E817K, H873P, R1105K, G1205E, R337stop, Q450stop, W937stop, and Q1434stop) to examine the functional implications of these variants (Pir et al., 2022; Pir et al., 2022). Furthermore, we employed the CRISPR system to introduce precise amino-acid substitutions at two different conserved amino acid positions (G680S and P702A) in *ift-140*, the ortholog of human IFT140 in *C. elegans*. Our functional analysis revealed that mutants bearing *ift-140*^W937stop^ or *ift-140*^P702A^ display null-mutant-like phenotypes, including lipophilic fluorescent dye uptake defect and IFT accumulations. Overall, our work shows that with the help of existing mutant resources and CRISPR/Cas9 systems, we can efficiently produce more data on human variants than we previously realized.

## Results

### Functional characterization of matching variants (MatchVars) in *C. elegans*

To leverage the already existing *C. elegans* mutants harboring variants, we initially gathered human and *C. elegans* variants for the *IFT140* gene from variant databases, including ClinVar, the Genome Aggregation Database (gnomAD), and the Trans-Omics for Precision Medicine (TOPMed), Wormbase and analyzed each position with ConVarT (Congruent clinical Variation Visualization Tool) to locate matching variants (MatchVar) between human IFT140 and *C. elegans* IFT-140 (**Figure 1 and Supplementary Table 1**). 80% of variation types in human *IFT140* presented by ClinVar are single nucleotide alterations, including 761 missense variants (13th February 2023) (**Figure 2A and B**). The functional classifications of these variants in human *IFT140* are far from complete because 80% of these missense variants are labeled as VUS (**Figure 2B**).

**Figure 1:**
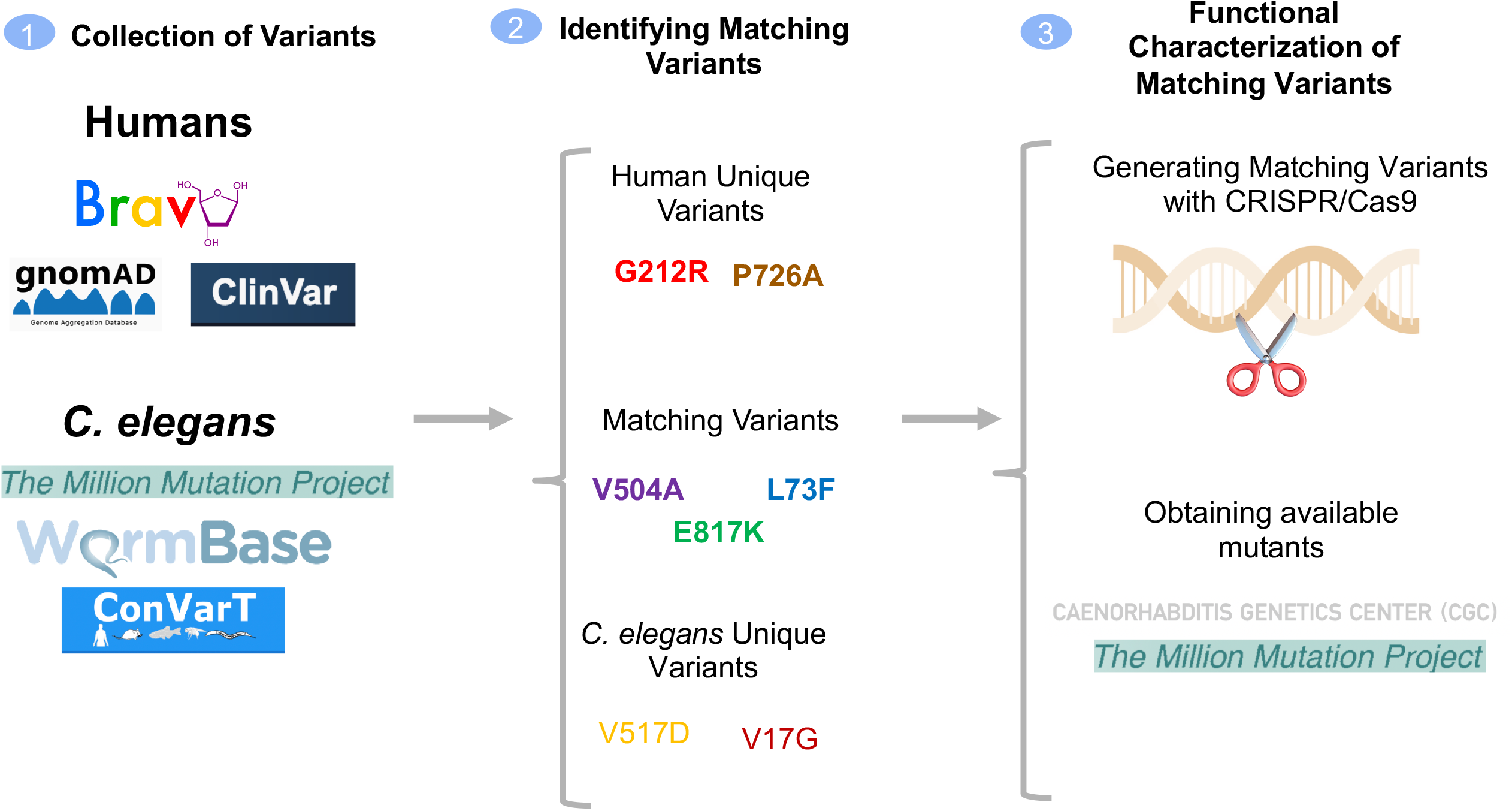
The workflow for variant assessment. The workflow of the current study is displayed. *C. elegans* and human variants for *IFT140* were gathered from the indicated resources along with relevant info, such as allele frequency and clinical significance. Following the discovery of matching (equivalent) variants (MatchVars) and distinct variants, mutants bearing a missense or a stop codon were obtained from the Caenorhabditis Genetics Center (CGC), or MatchVars were produced using the CRISPR/Cas9 genome editing tool.

**Figure 2:**
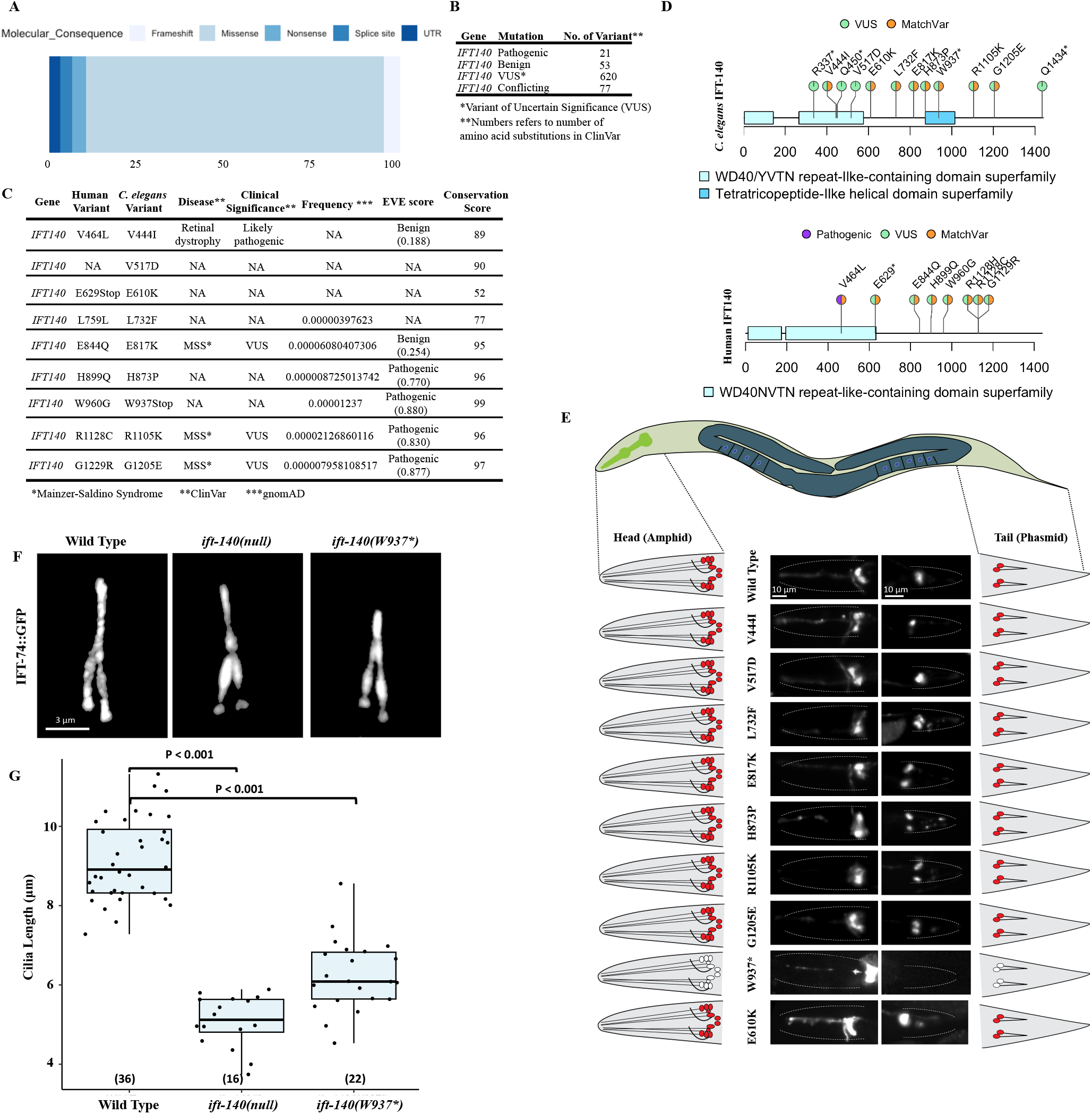
The *ift-140*^W937stop^ mutants phenocopy defects of loss-of function *ift-140* mutants. **A)** Shown are the percentage distribution of the variation types for human *IFT140* in the ClinVar. **B)** Number of missense variants in human *IFT140* from ClinVar is displayed with clinical significance. **C)** *C. elegans* variants in *IFT-140* and human variants in *IFT140* corresponding to the human positions of *C. elegans* variants are presented. Clinical significance, disease and frequency (downloaded from gnomAD), EVE score, and conservation score (see materials and methods) were presented for human variants. **D)** The variants, including matching variants (MatchVars) from humans and *C. elegans* IFT140 were displayed in their corresponding positions. VUS means variant of uncertain significance. **E)** Fluorescence images from the head and tail were displayed with representative drawings following the dye-filling assay. *C. elegans* cartoon shows the whole worm. The intermittent line in fluorescence images marks the edge of worms in the head and tail. With exception of mutants carrying W937stop, wild type and other worms absorb the lipophilic fluorescent dye in the head and tail. Asterisk (*) refers to the stop codon. Scale bar: 10 μm. **F)** Shown are the localization of IFT-74::GFP in the tail cilia (phasmid) of wild type, *ift-140(lf)*, and *ift-140*^W937stop^. Scale bar of 3 μm. **G)** The average tail cilia lengths for wild type, *ift-140(lf)*, and *ift-140*^W937stop^ were presented. The number of measured cilia was displayed in parentheses.

Our analysis revealed that *C. elegans ift-140* mutants contain a total of 39 variants (Pir et al., 2022; Pir et al., 2022). However, we restricted our analysis to 12 mutants, primarily focusing on MatchVars and as well as stop codons. Because the stop codon-carrying mutants would reveal which domains are important for the function of IFT-140. Human variants in IFT140 in the same position as those in *C. elegans* were listed together with the disease association, clinical significance, allele frequency, Evolutionary model of Variant Effect score (EVE), and conservation score for each variation (**Figure 2C**) (Frazer et al., 2021; Karczewski et al., 2020). ConVarT indicates that multiple variants, including *C. elegans* R1105K > human R1128C and *C. elegans* G1205E > human G1229R, are likely MatchVars (**Figure 2D**). Both R1128C and G1229R in human IFT140 are Mainzer-Saldino Syndrome-related variations of unknown significance (VUS). We obtained all these 12 mutants and a null *ift-140* mutant from Caenorhabditis Genetics Center (CGC) (**Supplementary Table 1**). We found that several mutants, including VC20793 (IFT-140(Q450stop)), had delayed development and excluded them from further investigation.

We next went on to perform a functional investigation of each mutant. The structural intactness of the cilia in *C. elegans* is frequently examined using the fluorescent lipophilic dye (DiO) uptake assay. The sensory neurons in the head (amphid) and tail (phasmid) of the wild type completely take in the dye via their cilia, whereas the failure of DiO uptake in the head (amphid) and tail (phasmid) frequently indicates structurally compromised cilia as observed with *ift-140* loss-of-function (lf) mutants (**Figure 2E**). We anticipate finding dye absorption failure if mutants exhibit structural ciliary defects comparable to the null *ift-14*0 mutant. Consistent with expectation, two different mutants (R337stop and W937stop), but not others display dye uptake failure, suggesting they might have defective cilia structures (**Figure 2E**). Furthermore, we performed the dye filling uptake assay following a shift to 25 °C for 16 hours to identify the temperature-sensitive mutants, and mutants bearing the V444I mutation exhibit a minor dye uptake deficiency, but further examination found no significant structural abnormalities in two distinct cilia (Data not shown). Both R337stop and W937stop mutants were independently crossed into CRISPR knock-in IFT-74::GFP allele and they display cilia IFT accumulations resembling the null *ift-14*0 mutant, but the dye uptake defect of R337stop mutants could not be rescued with a functional IFT-140::GFP transgene, we, therefore, did not include the R337stop for further analysis and our predictions (**Figure 2F** and data not shown). Similar to *ift-140(lf)*, our confocal microscopy analysis demonstrates that *ift-140*^W937stop^ mutants are shorter than those of the wild type (**Figure 2G**). Taken together, our strategy reveals W937stop (removing the last 599 amino acids and domain) is likely a null allele of *ift-140*, but *ift-140*^V444I^, *ift-140*^V517D^, *ift-140*^E610K^, *ift-140*^L732F^, *ift-140*^E817K^, *ift-140*^H873P^, *ift-140*^R1105K^, and *ift-140*^G1205E^ variants are not phenotypic, meaning that they are more likely benign.

### P726A variant associated with short-rib thoracic dysplasia, but not G704S in IFT-140, is a loss of function mutation and leads to the ciliary accumulations of the intraflagellar transport

As a part of our ongoing efforts to investigate the functional impact of MatchVars, we chose two human variants in IFT140, including p.Pro726Ala and p.Gly704Ser (GenBank: NM_014714.3), and then employed the CRISPR/Cas9 technology specifically to introduce the corresponding amino acid substitutions [Human IFT140(P726A) → *C. elegans* IFT-140(P702A) and human IFT140(G704S) → *C. elegans* IFT-140 (G680S)] into the *C. elegans* IFT-140 (**Figure 3A**). The human G704S in the *IFT140* gene has not been implicated in *IFT140*-related disease whereas G212R and P726A (*C. elegans* P702A) compound heterozygosity in *IFT140* resulted in short-rib thoracic dysplasia in a patient, therefore P726A is probably pathogenic, despite there has not yet been any functional data to classify the P726A variant in the *IFT140* gene as a pathogenic variant (**Figure 3A**) (Forbes et al., 2018). Additionally, the ClinVar put the P726A variant in the *IFT140* gene in the category of conflicting pathogenicity interpretations. The position of P726 and G704 are highly conserved among IFT140 orthologs from the 1088 different species (Conservation scores: 94 for G704 and 97 for P726; **Figure 3B and Supplementary Fasta**). The *C. elegans* IFT140(G704S) and IFT140(P702A) are MatchVars of human IFT140(G704S) (dbSNP: rs150745099) and IFT140(P726A) (dbSNP: rs1057518064), respectively (**Figure 3C**). We, therefore, employed *C. elegans* to experimentally investigate the functional effects of these two MatchVars: G680S (human G704S) and P702A (human P726A) in *C. elegans*.

**Figure 3:**
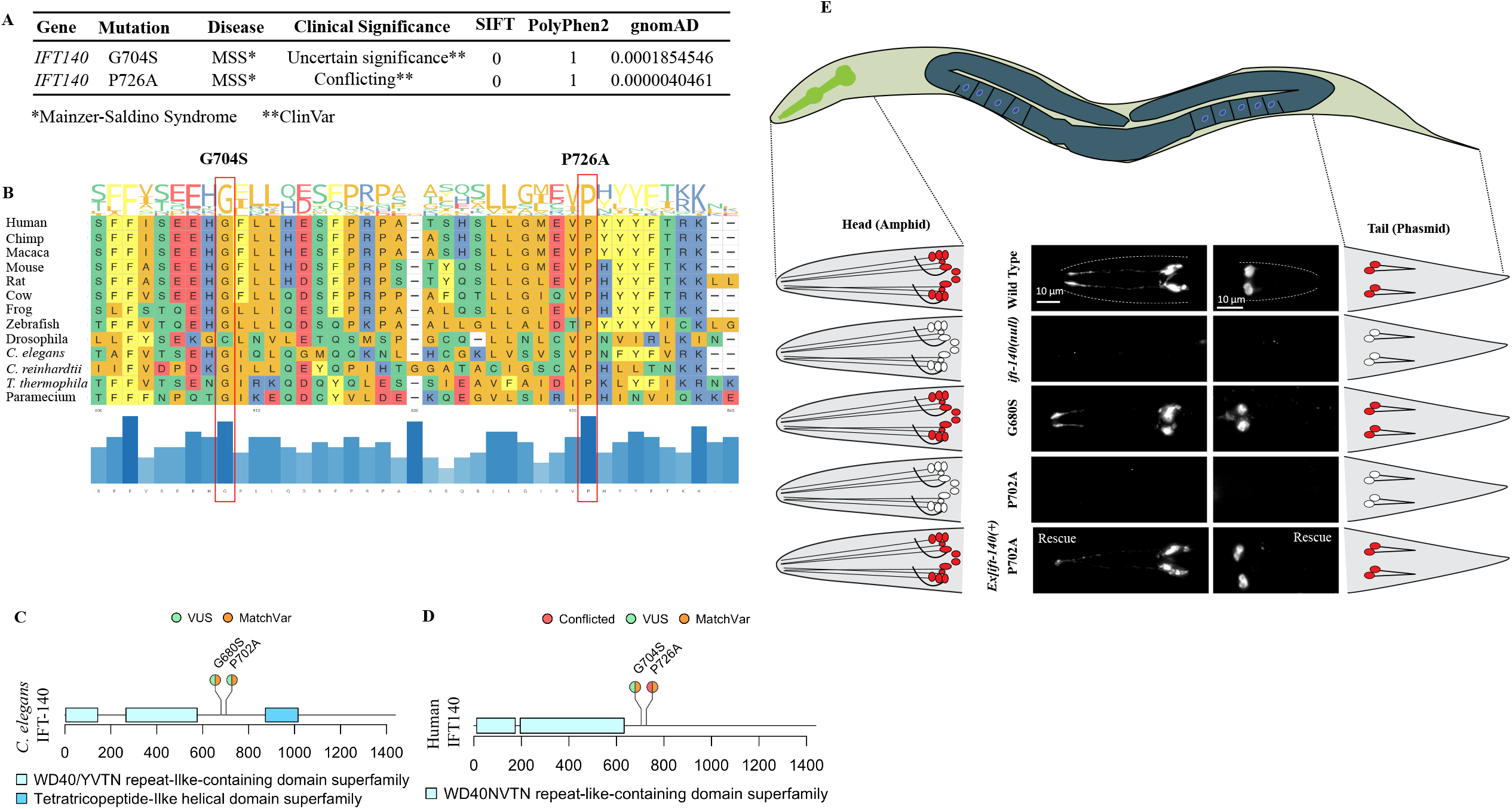
The *C. elegans* MatchVars for human IFT140^P726A^ but not IFT140^G704S^ displays defects in the lipophilic fluorescent dye uptake in the head and tail. **A)** Clinical significance and disease relevance from ClinVar, frequency from gnomAD, SIFT, and PolyPhen2 score were presented for two human variants, P726A and G704S. MSS stands for Mainzer-Saldino Syndrome. **B)** The positions of these two variants were shown on the MSA of IFT140 orthologs from 13 different species. The conservation of the corresponding amino acid position across the 13 species is depicted by the blue bars at the bottom of the plot. The extended versions of MSAs were provided as supplementary files. **C)** The positions of human P726A and G704S variants and *C. elegans* P702A and G680S variants were shown in a lollipop plot of IFT140 proteins from humans and *C. elegans*, respectively. The P702A and G680S are MatchVars of the human P726A and G704S in human IFT140, respectively. Clinical significance (VUS and conflicted) for human variants from ClinVar were presented. **D)** The entire worms, including heads and tails, were depicted in the *C. elegans* representative drawing. The fluorescence images were displayed alongside representative sketches of the head and tail cells. White indicates no dye uptake in the head and tail, while red indicates dye uptake in the cells. After the fluorescent dye assay, fluorescent images of the wild type and the indicated mutants were shown. *ift-140(lf)*, and *ift-140*^P702A^ mutants were Dye negative in the head and tails. The expression of *ift-140* completely restored the dye uptake failure of *ift-140*^P702A^ mutants. Scale bar: 10 μm

The DiO uptake assay results show that *ift-140(lf)* mutants and mutants carrying homozygous P702A but not homozygous G680S fail to take up the fluorescent dye, suggesting the P to A change at the position of 702 likely results in defects in cilia structure, which is a strong indicator of disruption of IFT140 function. Importantly, dye filling defect (Dyf) of *ift-140*^P702A^ mutants was fully rescued by introducing a wild type copy of *ift-140/IFT140* into homozygous *ift-140*^P702A^ mutants, suggesting Dyf is due to P702A variant in IFT-140 (**Figure 3D**).

IFT-140 is a critical part of IFT-A, hence its absence should cause IFT to be defective. We, therefore, crossed GFP-tagged IFT markers (endogenous IFT-74::GFP and OSM-3/KIF17::GFP) into *ift-140(lf), ift-140*^P702A^ and *ift-140*^G680S^ mutants to visualize the IFT. Our confocal microscopy analysis revealed that both *ift-140(lf)* and *ift-140*^P702A^ mutants but not *ift-140*^G680S^ display remarkably similar phenotypes with the ciliary accumulation of a GFP-tagged IFT core machinery components (**Figure 4A**). Furthermore, the PHA/PHB cilia are shorter in both *ift-140(lf)* and *ift-140*^P702A^ as compared to wild type and *ift-140*^G680S^ (**Figure 4B**). We next went on investigating other cilia types, including AWA and AWB. The AWA olfactory neurons possess complex cilia with multiple branches, whereas the AWB cilia are Y-type cilia, with short and long branches. Expectedly, the complexity of AWA drastically decreases in both *ift-140(lf)* and *ift-140*^P702A^ mutants, and the AWB cilia in both mutants display dramatically altered Y-type shape (**Figure 4C**).

**Figure 4:**
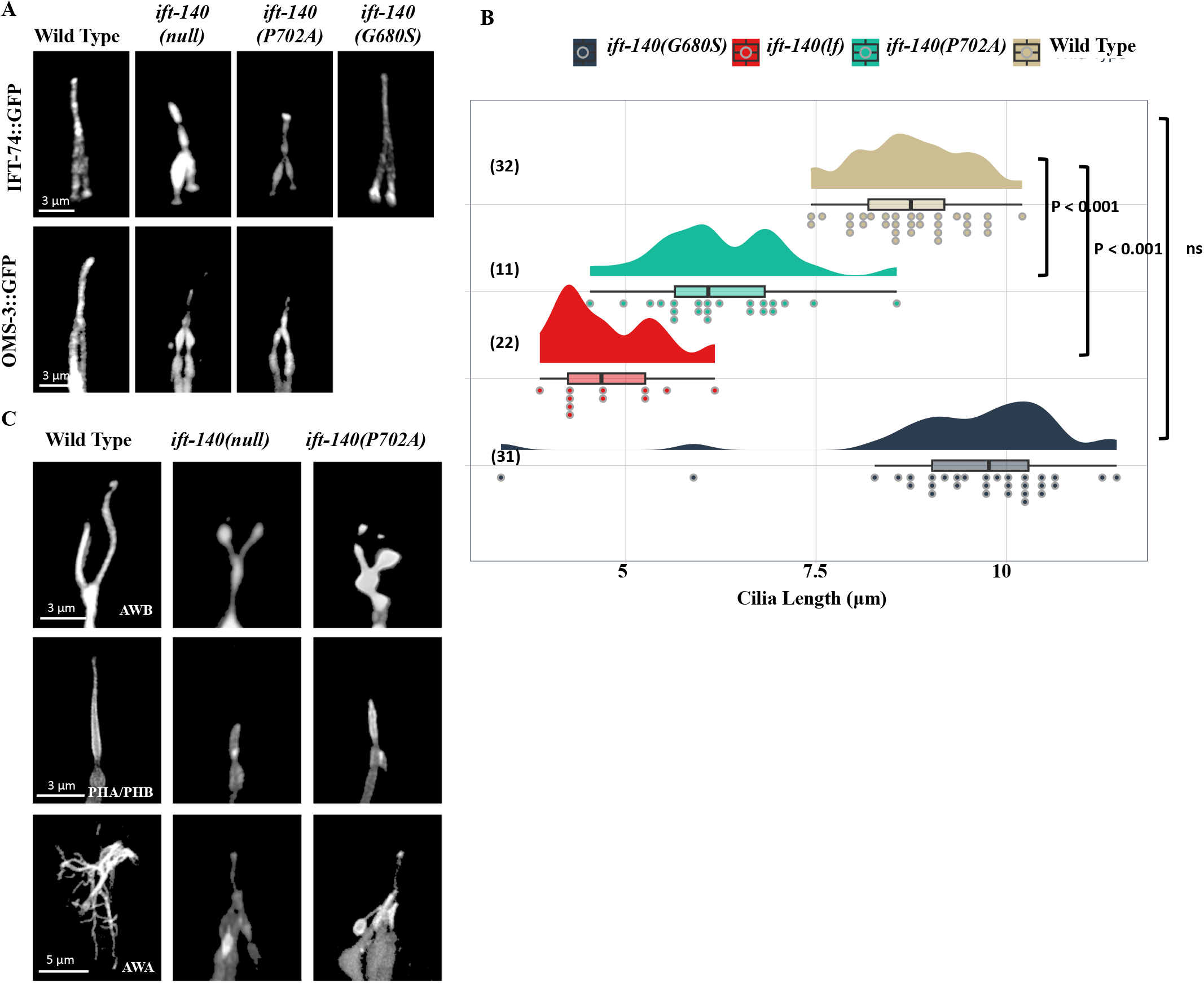
The sensory cilia in the *C. elegans* IFT-140^P702A^ mutants are shortened. **A)** Shown are confocal images displaying the localization of fluorescently tagged IFT proteins in the tail (phasmid) of wild-type and the indicated mutants. Scale bar: 3 μm. **B)** Plots of the PHA/PHB cilia length for wild-type and indicated mutants were shown. Not Significant is abbreviated as ns. P values between the designated mutants and the wild type are also presented. Numbers in parentheses indicate the number of cilia used to calculate cilia length. **C)** Fluorescent markers display the following cilia: AWB cilia (Y-shaped cilia), PHA/PHB cilia (rod-shaped cilia), and AWA cilia (multiple complex branches). Shown are representative fluorescent images from wild type, *ift-140(lf)*, and *ift-140*^P702A^ mutants. Scale bar (AWB): 3 μm, scale bar (PHA/PHB): 3 μm, and scale bar (AWA): 5 μm.

Both GFP-tagged odorant response abnormal protein 10 (ODR-10::GFP, AWA cilia) and OSM-9::GFP (QLQ cilia, *C. elegans* ortholog of human TRPV4) were independently crossed into both *ift-140(lf)* and *ift-140*^P702A^ mutants, and following confirmation, wild type and mutants expressing ODR-10::GFP or OSM-9::GFP were imaged using the confocal microscopy. Microscopy analysis confirms that AWA cilia are severely impacted in both *ift-140(lf)* and *ift-140*^P702A^ mutants. The ODR-10 protein was very low in the cilia in both mutants, but it was a bit difficult to determine whether this was due to significantly altered cilia structure or a decrease in ODR-10 staining in cilia (**Figure 5A**). However, OSM-9::GFP stains the whole QLQ cilia in both mutants, indicating that OSM-9::GFP enters cilia, and both mutants exhibit OSM-9 accumulations at the base of the cilia as compared to the wild type (**Figure 5A**). Our analysis reveals that both mutants appear to have altered QLQ cilia morphology. Taken together, *ift-140*^P702A^ affects the localization of membrane proteins.

**Figure 5:**
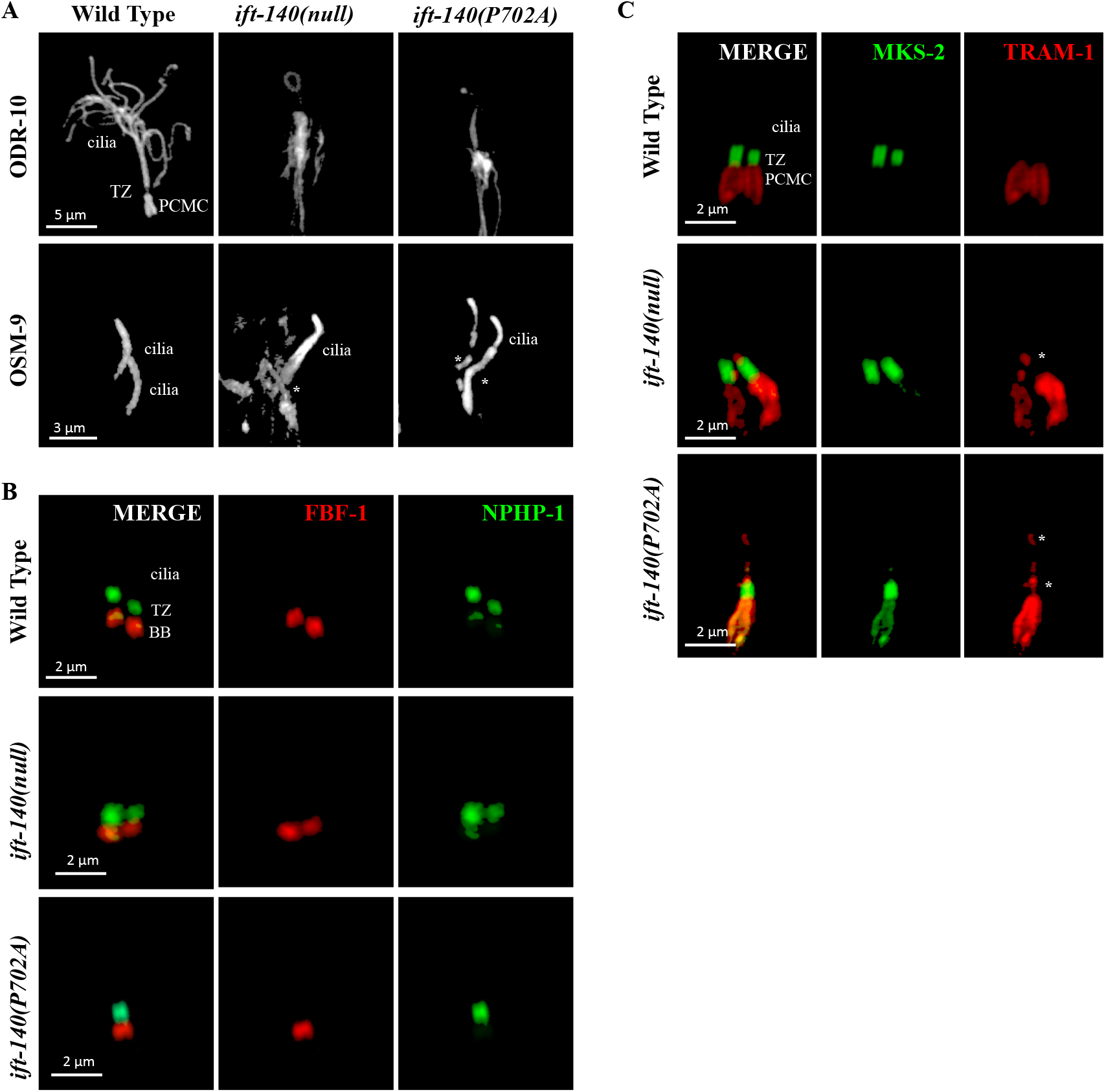
The ciliary gate is altered in *C. elegans* IFT-140^P702A^ mutants. **A)** Localization of fluorescently tagged membrane proteins (ODR-10 and OMS-9) were shown in wild type, *ift-140(lf)*, and *ift-140*^P702A^ mutants. Scale bar (ODR-10): 5 μm and scale bar (OSM-9): 3 μm. Asterisk (*) shows accumulation of OSM-9 at the base of cilia. **B)** Confocal images showing both the red-tagged DYF-19 (FBF-1, basal body protein) and the GFP-tagged NPHP-1 (a transition zone protein) were presented in wild type, *ift-140(lf)*, and *ift-140*^P702A^ mutants. BB and TZ stand for basal body and transition zone, respectively. Scale bar: 2 μm. **C)** Shown are confocal images from non-ciliary transmembrane protein TRAM-1 (labeled with red) and MKS-2 (green, a transition zone protein) in wild type, *ift-140(lf)*, and *ift-140*^P702A^ mutants. Asterisk (*) indicates the ciliary entry of TRAM-1. Scale bar: 2 μm.

A previous study revealed that IFT140 regulates the ciliary gate, thus we investigated the impact of P702A variant on the ciliary gate and protein trafficking (Scheidel and Blacque, 2018). TRAM-1 (Translocation Associated Membrane Protein 1 orthologue) was not inside cilia in the wild type, but TRAM-1 enters cilia in both *ift-140(lf)* and *ift-140*^P702^ mutants whereas the localization of transition zone proteins (NPHP-1; nephrocystin-1 orthologue) remains unaltered in these mutants (**Figure 5B and C**). Taken together, multiple lines of evidence suggests that *C elegans* MatchVars for human IFT140^P726A^ but not IFT140^G704S^ disrupts the functions of *IFT140*, and *C elegans ift-140*^P702A^ change likely represents a loss of function variant of *IFT-140*.

## Discussion

Determining the consequence of human variants is important for diagnosing genetic diseases, and for guiding the development of drug response regimes. It has been widely accepted that analyzing MatchVars from model organisms might shed light on the consequences of human MatchVars, including understanding the association between human variants and diseases (AlAbdi et al., 2022; Di Rocco et al., 2022; Lange et al., 2022; Macaisne et al., 2022; Morbidoni et al., 2021; Platzer et al., 2019; The Alliance of Genome Resources Consortium et al., 2020; Wang et al., 2022, 2017; Zhu et al., 2020). Recently, we published ConVarT, a search engine that displays disease and phenotypic data related to variants, including MatchVars from humans, mice, and *C. elegans* on multiple sequence alignments of orthologous genes from these three species (Pir et al., 2022; Pir et al., 2022). The Australian Phenome Bank (APB) is a database of mouse strains carrying variations, including MatchVars in different genes, while the Million Mutation Project generated 2000 *C. elegans* mutants with many missense mutations in various genes (Thompson et al., 2013). 2359 mice mutant strains (5th February 2023) are currently available through APB. The capability to explore the consequence of a single missense variant offers a great advantage for modeling the corresponding human MatchVars. Thus, these available *C. elegans* and mice mutant databases are valuable resources when searching for MatchVars for functional analysis. Indeed, we take advantage of the availability of *C. elegans* mutants bearing MatchVars and demonstrate that the use of these mutants can provide experimental evidence for the classification of human MatchVars. Our findings support the assertion that the MatchVars are useful tools for systematically assessing the molecular consequence of human variants and they can help provide functional proof for the clinically relevant variants. Moreover, CRISPR-mediated generation of MatchVars contributes to increasing the pools of MatchVars for functional evaluations, and the *C. elegans* community continuously creates MatchVars to evaluate their magnitude. This not only contributes to the functional characterization of human MatchVars in *C. elegans* but also provides valuable and independent resources for clinical scientists. This can be complementary to other findings for human variants. For example, the availability of phenotypic MatchVars from model organisms can be a great resource for clinical scientists when they look for evidence for a variant to decide on classifying it as a potential disease-causing variant.

In the current study, we chose *ift-140* for further evaluation because the phenotypic characterization of *ift-140* null mutants is straightforward. We first used ConVartT (https://convart.org/), which provided multiple sequence alignments along with the human, mouse, and *C. elegans* variants, to identify variants of interest (Pir et al., 2022; Pir et al., 2022). We focused on 12 mutants, primarily studying MatchVars, and added two additional MatchVars using the CRISPR/Cas9 system. Expectedly, the Q1434stop variant of IFT140 (NCBI Reference Sequence: NP 506047, and 1437 amino acids) did not result in any anomalies in cilia because it just deletes the last three amino acids (data not shown). However, removing the last 500 amino acids (W937stop) containing half of the tetratricopeptide repeat (TPR) causes ciliary IFT accumulation, with short cilia, indicating the functional importance of a tetratricopeptide-like helical domain. Consistent with this view, a recent study revealed that the TPR domain of IFT140 is important for its interaction with the TPR domains of IFT144 (Hesketh et al., 2022).

Our *C. elegans* work experimentally measured the functional consequence of eight missense variants in *C. elegans* IFT-140 (V444I, V517D, E610K, L732F, E817K, H873P, R1105K, and G1205E). Before our study, all of these mutations were thought to be VUS; nevertheless, our findings classify them as benign. Interestingly, the evolutionary conservation scores from multiple sequence alignments of 1088 IFT140 orthologs revealed that several residues (V517, E817, H873, R1105, and G1205) are highly evolutionarily conserved positions (conservation score ≥ 90) (**Supplementary Fasta**). However, none of these variations are situated in the amino acid position necessary for interaction between human IFT140 and IFT144 as the recent study revealed that in the human IFT140, many positions, including D at position 789, F at position 792, K at position 796, V at position 822, N at position 826, A at position 830, A at position 833, and R at position 837, are crucial for the expected interface between IFT140 and IFT144 (Hesketh et al., 2022).

The V464L missense variant in human IFT140 (dbSNP: rs2034681207 and NP 055529.2) was submitted to ClinVar as a likely pathogenic for Retinal dystrophy, however, there is no functional evidence to support this claim. Our functional analysis suggests the V444I missense variant in *C. elegans* IFT-140 to be a benign mutation. Does the functional data with the V444I missense accurately reflect the V464L missense variant in the human IFT140 gene because the alteration is not the same? It could be due to the nonpolar and hydrophobic nature of the amino acids leucine (Leu) and isoleucine (Ile). Furthermore, consistent with our suggestion, the EVE (evolutionary model of variant effect) online tool predicts that the V464L missense variant in human IFT140 is likely benign (Frazer et al., 2021). In addition, the CRISPR/Cas9 system produced two missense variants (P702A and G680S) in IFT-140 in *C. elegans*, which are the MatchVars of P726A and G704S in human IFT140, respectively. In contrast to human IFT140^G704S^, the MatchVar of the human IFT140^P726A^ missense variant represents a loss of IFT-140 function in *C. elegans*. The human IFT140^P726A^ missense variant is a conflicting variant for clinical significance, and our findings provide functional evidence for the variant in favor of pathogenicity for the first time. Taken together by employing available mutant resources from model organisms and CRISPR/Cas9 systems in model organisms in determining the consequence of MatchVars, we can systematically generate more knowledge about human variants than previously realized.

## Materials and Methods

### Fluorescent lipophilic dye uptake assay

The fluorescent lipophilic dye was diluted with M9 (1:200 fluorescent lipophilic dye: M9 and fluorescent lipophilic dye were stored at -20°C) and incubated with mixed stages of healthy, well-fed *C. elegans* worms for 45 minutes. Wild types were always included in each Dye assay, while contaminated plates (bacteria or fungi) were not examined. At least 50 worms were scanned for each independent experiment. After it was confirmed that wild types fully absorbed Dye, microscope images were collected. Following the *ift-140(syb1325)V*.*(P702A);Ex[IFT-140::GFP+pRF4]* dye uptake assay, the dye rescue assay plate containing mutants with or without *Ex[IFT-140::GFP+pRF4]* was photographed to ensure that the dye uptake is fully restored. The same procedure was carried out for the plate carrying the mutation *che-11(gk925030) (R337stop);Ex[IFT-140::GFP+pRF4]*, however there was no rescue for the dye uptake defect of *che-11(gk925030) (R337stop)*. To better understand the cause of the dye uptake defect failure, *IFT-140::GFP* in *che-11(gk925030) (R337stop)* were photographed, which showed the ciliary accumulations of *IFT-140::GFP*.

### Confocal microscopy imaging and subsequent image analysis

Before beginning the microscope analysis, 1 μl of 10 mM levamisole was applied to the freshly prepared microscope slides with a 2-3 percent agarose pad. These slides were placed into the microscope. Before the microscopy imaging, the worms were checked to determine whether worms were well-fed and healthy. Next, microscope images were collected with the Zeiss LSM900 confocal microscope equipped with Airyscan 2 and controlled by ZEN 3 Blue edition software. A Plan ApoChromat 63x/1.40 NA with 0.14 μm intervals was used to collect Z-stack images, and then the Blue edition software ZEN 3 generated Z-stack generation. ImageJ (NIH) and Fiji software were used for the subsequent image analysis (Schindelin et al., 2012).

### Strains and Genetic Cross

The strains are maintained in a 20 °C incubator on standard Nematode Growth Medium (NGM) plates and fed with *E*.*coli* OP50 bacteria (Brenner, 1974). Standard crossing techniques were utilized to generate transgenic worms with various mutant backgrounds. The rescue lines were generated by crossing *Ex[IFT-140::GFP+pRF4]* into the *ift-140(P702A)* mutant. PCR and dye filling assay were used for confirmation of genotypes (genotyping *ift-140(P702A)* are available in **Supplementary Figure 1**). *Ex[IFT-140::GFP+pRF4]* were crossed into VC41005 [(ift-140 (gk925030) (R337stop)] mutants, the dye assay were used for confirmation. For the complete mutants they produced, the Million Mutation Project did whole genome sequencing (WGS) for VC40309[(ift-140(gk566606) (W937stop)] and VC41005[(ift-140(gk925030) (R337stop)] revealed the details of mutations. The presence of mutations in the VC40309[(che-11(gk566606) (W937stop)] and VC41005[(ift-140(gk925030) (R337stop)] mutants was confirmed by Sanger sequencing. All strains and primers generated by the CRISPR/Cas9 system and genetic crossing were listed in **Supplementary Table 4** and **Supplementary Table 5**.

### CRISPR/Cas9 mediated generation of variants in *IFT-140 in C. elegans*

The Bristol N2 strain was used as the wild type background strain for CRISPR/Cas9 editing experiments. Genome editing was conducted as previously described but with small modifications (Arribere et al., 2014; Kim et al., 2014). In brief, sgRNA plasmid, Cas9 plasmid, reporter marker plasmid, repair template (oligo or plasmid), Co-CRISPR plasmid or Co-conversion sgRNA plasmid (and Co-conversion repair oligo) were injected into N2 animals. F1 Animals with Co-CRISPR or Co-conversion phenotype were isolated and cultured in a single Nematode Growth Medium (NGM) plate. The F1 animals were lysed for screening of heterozygotes with targeted gene editing. F2 animals were cloned for homozygotes screening. The homozygotes were verified by PCR amplification and sequencing. The sgRNA target sites used in this study are listed in **Supplementary Table 2**. For the repair templates, we used the long oligos in IFT-140(G680S) editing and plasmid in IFT-140(P702A) editing. For the oligo templates, the homology arm is about 50 nt on either side of the mutated site. For the plasmid template, the homology arm is about 400bp on either side of the mutated site. The nucleotide mutations generated in each gene editing experiment are listed in the **Supplementary Table 3**. In IFT-140(P702A) editing experiments, precise gene editing introduced a restriction enzyme (Nru I) site. We thus amplify the genomic DNA sequences spanning each of the above four sites, and screen for the mutated animals through corresponding restriction enzyme digestion. For the IFT-140(G680S) editing, as there is no restriction site generated or destroyed, we used the allele-specific primers to screen for heterozygotes. The sequencing primers and genotyping primers used in each gene editing experiment are listed in the **Supplementary Table 4**.

### Multiple Sequence Alignments, Conservation Score, Plot Generation, and Statistical Analysis

Following the input of human IFT140 protein sequences (Human-NP_055529) into protein-protein Blast (BlastP), the following settings were used: Max target sequences :5000, Max target sequences :0.05, Matrix: BLOSUM62. 3232 proteins were displayed in the BlastP findings, and FAST complete sequences were obtained from NCBI. Eventually, the query covers less than 40, and proteins from the same organisms were removed from the download file to eliminate duplicated and incorrect orthologs. Using the msa R package (packageVersion: ‘1.30.1’), multiple sequence alignments (MSA) of IFT140 orthologs from 1088 organisms were carried out to visualize the conservation score for each position (Bodenhofer et al., 2015; Winter, 2017). For a representative MSA and plot, the following organisms were chosen and the RefSeq Protein IDs of the following organisms were gathered: Human-NP_055529, Chimp-XP_016784643, Macaca-XP_001089057, Cow-XP_002697959, Mouse-NP_598887, Rat-XP_006246116, Frog-NP_001116497, Zebrafish-XP_695732, Fruit Fly-NP_995608, *C. elegans*-NP_506047, *C. reinhardtii*-XP_042921850, *T. thermophila*-XP_001020653, and *Paramecium tetraurelia*-XP_001347023. The rentrez package was used to download the protein sequences of these selected organisms after creating tables in R with accession numbers. MSA was carried out with the msa R package (Bodenhofer et al., 2015; Winter, 2017). Plots were generated using the following R packages, including trackViewer (packageVersion: ‘1.34.0’), ggmsa (packageVersion: ‘1.4.0’), and Welch’s two-sample t-test (statistical analysis) were performed using R (Ou and Zhu, 2019; Zhou et al., 2022). R 4.2.2 was used throughout the R analysis.

### Collection of ClinVar variants

IFT140 input was placed into the ClinVar website (https://www.ncbi.nlm.nih.gov/clinvar/). The numbers on the bottom left of the screen displayed the variation type, molecular consequences, and clinical significance. The variation type was plotted in Figure 2A. Selected missenses with molecular consequences were displayed in Figure 2B.

## Author contributions

PZ and WB generated two mutants used in Figures 3, 4, and 5. AZ generated many strains, but due to her pregnancy and the subsequent delivery of her gorgeous child, she was abruptly forced to leave her work. Many congrats! Sadly, all of her strains—with the exception of one—were discovered to be lost and not frozen. The project was taken over by SC, who expanded it by creating all missing strains and additional strains to complete the work. SC carried out the all microscopy analyses. SC and OIK designed, and supervised the project, and prepared the figures. OIK wrote the paper.

## Acknowledgment

We thank Furkan Torun and Mustafa S. Pir for helping to initiate the project, and Furkan Kepenek for MSA. Although we would love to thank a funding agency for their support, we regrettably did not receive any financial support for this project. Thus, we collaborated with SunyBiotech. Lastly, we would be happy to share all reagents generated as a part of this study.

